# Depolymerized lamins link nuclear envelope breakdown to mitotic transcriptional quiescence

**DOI:** 10.1101/334110

**Authors:** Kohta Ikegami, Stefano Secchia, Jason D. Lieb, Ivan P. Moskowitz

**Affiliations:** Departments of Pediatrics, The University of Chicago, Chicago, Illinois, USA; Departments of Pathology, The University of Chicago, Chicago, Illinois, USA; Departments of Human Genetics, The University of Chicago, Chicago, Illinois, USA; Department of Biology, Lund University, Lund, Sweden

## Abstract

The nuclear envelope, a defining feature of eukaryotic cells, restricts DNA-dependent processes including gene transcription to the nucleus. The nuclear lamina is an integral component of the animal nuclear envelope, composed of polymers of nuclear lamin proteins^1,2^. Upon mitosis, the nuclear lamina disassembles, the nuclear envelope breaks down, and transcription becomes quiescent^3,4^. We report here a direct molecular link between nuclear lamina disassembly and mitotic transcriptional quiescence. We found that, at the G2 cell-cycle phase immediately preceding mitosis, nuclear lamin A/C (LMNA) became phosphorylated at Ser22 and depolymerized from the nuclear lamina. Depolymerized LMNA accumulated in the nuclear interior and physically associated with active *cis*-regulatory elements genome-wide. Depolymerized LMNA-associated sites were overrepresented near genes repressed by LMNA, suggesting that depolymerized LMNA participates in transcriptional repression at G2. Consistently, depolymerized LMNA-target genes underwent a steep expression decline from S to G2/M. Furthermore, *LMNA* deletion caused inappropriate RNA Polymerase II (Pol II) accumulation downstream of Pol II pause sites at promoters and enhancers genome-wide, leading to inappropriate and excessive transcriptional elongation. A subset of depolymerized LMNA-target genes were upregulated in fibroblasts of patients with Hutchinson-Gilford progeria, a premature aging disorder caused by *LMNA* mutations^5^, raising the possibility that defects in depolymerized LMNA-mediated mitotic transcriptional quiescence contribute to progeria pathogenesis. These observations support a model in which depolymerized LMNA targets active regulatory elements to promote RNA Pol II pausing preceding mitosis, coupling nuclear envelope breakdown to mitotic transcriptional quiescence.

## MAIN

LMNA serves as a constituent of the polymers that form the nuclear lamina, a protein meshwork that underlies the inner nuclear membrane^2^. LMNA polymers contribute to chromosome organization by interacting with megabase-wide regions of transcriptionally inactive domains in the genome called lamina-associated domains (LADs)^6–11^. During mitosis, LMNA is in the depolymerized form^4^. LMNA depolymerization is initiated by phosphorylation at Ser22 (S22) and Ser392 (S392), which is mediated by CDK1-Cyclin B, a kinase complex that promotes the cellular transition from G2 to mitosis^4,12–14^ (**Fig. 1a**). Previous studies show that LMNA depolymerization precedes the nuclear envelope breakdown in mitosis^15^ and that S22-phosphorylated LMNA is detectable prior to mitosis^16^. To define the precise cell-cycle time at which LMNA depolymerizes, we performed immunofluorescence microscopy for S22-phosphorylated (pS22) LMNA in the hTERT-immortalized human fibroblast cell line BJ-5ta (**Fig. 1b**). We found that pS22-LMNA became highly abundant in the nuclear interior in G2-phase cells (“G2” in **Fig. 1c**) before the onset of chromatin condensation (“Late G2” in **Fig. 1c**), before the appearance of prophase marker phospho-H3S10^17^ (**Fig. 1d**), and before nuclear envelope breakdown (“prometaphase” in **Fig. 1c**). The accumulation of pS22-LMNA was concomitant with progressive loss of the unphosphorylated LMNA signal at the nuclear periphery (“G2”, “Late G2”, and “prometaphase” in **Fig. 1c**), consistent with the paradigm that pS22-LMNA is generated by depolymerization of nuclear-peripheral LMNA.

**Figure 1.**
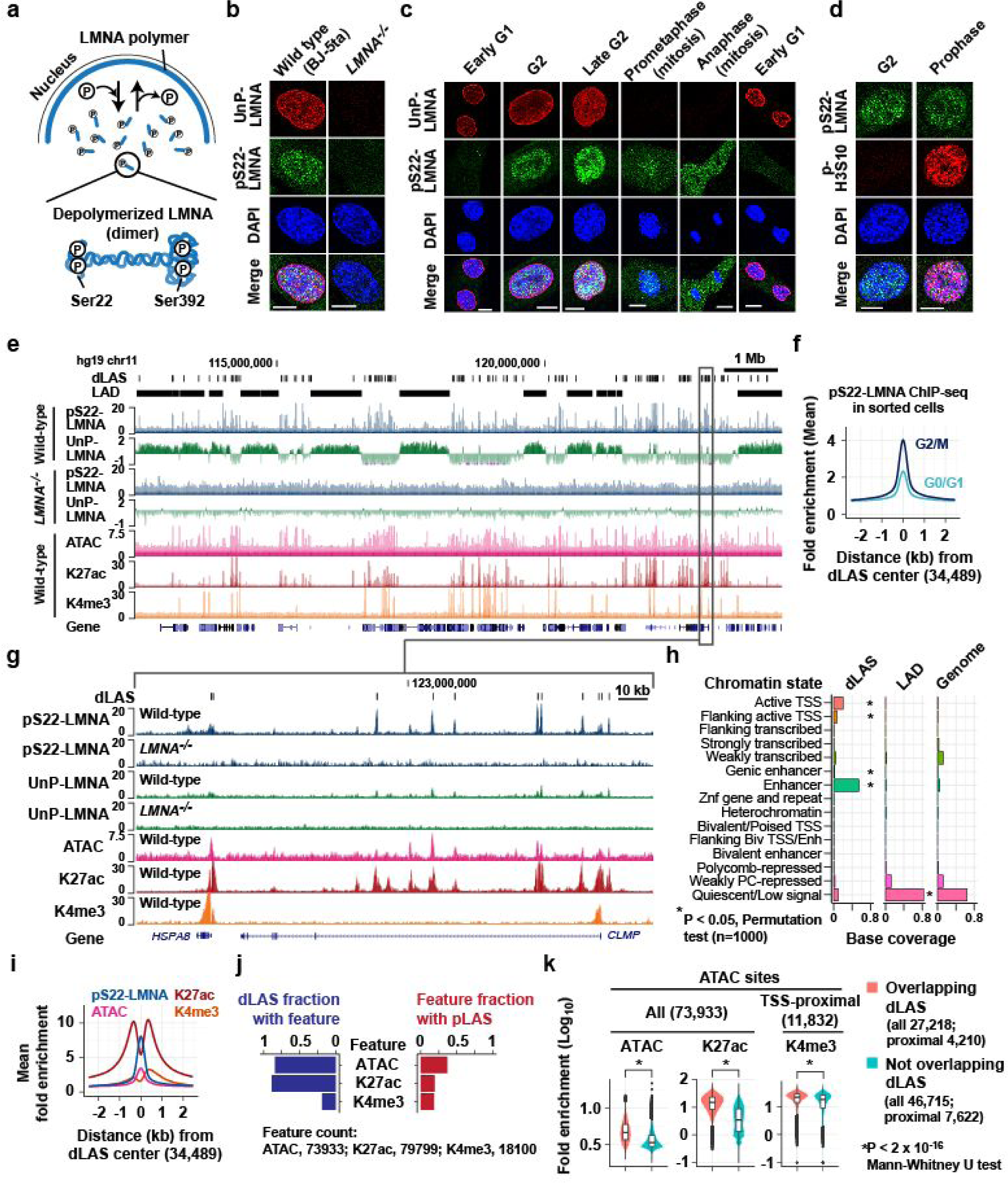
Depolymerized LMNA associates with active regulatory elements genome-wide. **a.** Depolymerized LMNA is phosphorylated at Ser22 and Ser392. **b.** Immunofluorescence for unphosphorylated (UnP-) and Ser22-phosphorylated (pS22-) LMNA in BJ-5ta human fibroblast cell line (wild type) and BJ-5ta-derived *LMNA*^*–/–*^ cell line. Bar, 10 µm. **c.** Same as **b**, but wild-type BJ-5ta cells at specific cell-cycle stages are shown. **d.** Immunofluorescence for pS22-LMNA and prophase marker phospho-H3S10 in BJ-5ta cells. **e.** A representative genomic region showing pS22-LMNA ChIP-seq, UnP-LMNA ChIP-seq, ATAC-seq, H3K27ac ChIP-seq, and H3K4me3 ChIP-seq signals in asynchronous BJ-5ta cells. UnP-LMNA profiles are in the log_2_ scale to show low-level enrichment characteristic of lamina association. dLAS, depolymerized LMNA-associated site. LAD, lamina-associated domain. **f.** pS22-LMNA ChIP-seq signals at dLASs in BJ-5ta cells sorted for the G2/M or G0/G1 cell-cycle stage. **g.** ChIP-seq and ATAC-seq profiles in the region indicated in **e**. **h.** dLASs and LADs location (defined in BJ-5ta) with respect to the 15-state chromatin annotations in normal dermal fibroblasts. **i.** ChIP-seq and ATAC-seq signals at dLASs in BJ-5ta. dLASs are oriented 5′ to 3′ relative to the closest gene orientation. **j.** (Left) Fraction of dLASs that overlap ATAC, K27ac, or K4me3 peaks. (Right) Fraction of ATAC, K27ac, or K4me3 peaks that overlap dLASs. **k.** ATAC-seq, K27ac, or K4me3 ChIP-seq signals at ATAC sites overlapping or not-overlapping dLASs. TSS-proximal ATAC sites are those located within –1kb to +500 bp of TSSs.

We sought to determine the localization of depolymerized LMNA in the G2 nuclei. To this end, we performed ChIP-seq in asynchronous BJ-5ta cells using the antibody specific to pS22-LMNA. In sharp contrast to LMNA polymers that interacted with megabase-wide LADs (**Fig. 1e; Extended Data Fig. 1a, c**), depolymerized LMNA exhibited point-source enrichment at discrete sites located outside LADs (**Fig. 1e; Extended Data Fig. 1a–d**). We identified 34,489 depolymerized-LMNA-associated sites (dLASs) genome-wide. The depolymerized LMNA ChIP-seq signals were completely abolished in a *LMNA*^*–/–*^ cell line derived from BJ-5ta (**Fig. 1e; Extended Data Fig. 1b**). Because depolymerized LMNA becomes abundant at G2 (**Fig. 1c**), we hypothesized that the dLAS ChIP signal from the asynchronous cell population emanated from cells in the G2 stage of the cell cycle. We reperformed pS22-LMNA ChIP-seq on cell-cycle sorted BJ-5ta cells, and observed strong enrichment of depolymerized LMNA at dLASs in G2/M relative to G1/G0 (**Fig. 1f**). To examine the chromatin localization of depolymerized LMNA with a different approach, we performed anti-Ty1 tag ChIP-seq on BJ-5ta cells overexpressing a Ty1-tagged phospho-deficient (S22A/S392A), or a phospho-mimetic (S22D/S392D) LMNA mutant. Phospho-mimetic Ty1-LMNA-S22D/S392D was strongly enriched at dLASs (**Extended Data Fig. 1e**), and 80% of its bound sites (31,392 sites) overlapped 73% of dLASs (**Extended Data Fig. 1f**) while phospho-deficient LMNA associated with only 6,044 sites, overlapping 18% of dLASs, and showed far weaker enrichment at dLASs (**Extended Data Fig. 1e, f**).

The vast majority of the 34,489 dLASs were located at accessible chromatin regions and marked by histone modifications characteristic of active regulatory elements (**Fig. 1g–i**): 84% of dLASs coincided with ATAC-seq-defined accessible chromatin regions; 80% with H3K27ac-marked putative active promoter/enhancer regions; and 17% with H3K4me3-marked putative active promoter regions in BJ-5ta (**Fig. 1j**). A comparison with the chromatin state annotations in normal dermal fibroblasts^18^ revealed that 14% of dLASs were marked as the “Active TSS” state (P < 0.05) and 54% as the “Enhancer” state (P < 0.05) (**Fig. 1h**). Although nearly all dLASs corresponded to accessible chromatin, dLASs marked a modest subset (37%) of all accessible chromatin sites in the genome (total 73,933 sites) (**Fig. 1j; Extended Data Fig. 1g**). These dLAS-associated subset of accessible chromatin regions exhibited higher levels of chromatin accessibility, H3K27ac, and H3K4me3 relative to the non-associated subset (**Fig. 1k**). Thus, depolymerized-LMNA associates with the highly active subset of gene-proximal and distal regulatory elements genome-wide.

Prompted by the localization of depolymerized LMNA at putative regulatory elements, we tested whether *LMNA* deletion affects transcription by profiling nascent RNAs using GRO-seq in asynchronous wild-type and *LMNA*^*–/–*^ cells. Among 11,248 protein-coding genes with detectable GRO-seq signals (GRO-seq RPKM > 0.1; termed “transcribed genes” hereafter), *LMNA* deletion resulted in increased transcription of 358 genes (termed “*LMNA*^-/-^-up genes”) and decreased transcription of 550 genes (termed “*LMNA*^-/-^-down genes”), together termed “*LMNA*-dependent genes” (**Fig. 2a**). *LMNA*-dependent genes were located predominantly outside LADs: Only 17 of 358 *LMNA*^-/-^-up genes (5%, P=0.03) and 27 of 550 *LMNA*^-/-^-down genes (5%, P=0.005) had TSSs within LADs (**Fig. 2b**). Furthermore, *LMNA*-dependent gene TSSs were not located significantly closer to LADs compared to all transcribed genes (P=0.18, *LMNA*^-/-^-up genes; P=0.31, *LMNA*^-/-^-down genes; **Fig. 2c**). In contrast, 91% of all transcribed genes contained at least one dLAS within +/-100 kb of their TSSs (**Fig. 2d**). Interestingly, dLASs were not evenly distributed among transcribed genes: *LMNA*^*–/–*^-up genes contained a large number of dLASs within +/-100 kb TSS relative to all transcribed genes (51% of *LMNA*^*–/–*^-up v. 37% of all transcribed genes for ≥5 dLASs, P=1.1×10^−8^; **Fig. 2d**), and were significantly closer to dLASs relative to all transcribed genes (P=1.7×10^−18^; **Fig. 2e**). Similarly, 102 genes upregulated by overexpression of the LMNA-phospho-deficient mutant (S22A/S392A) extensively overlapped *LMNA*^*–/–*^-up genes (31%, P=1.3×10^−23^; **Extended Data Fig. 2**) and showed high densities of dLASs relative to all transcribed genes (**Extended Data Fig. 3**). In contrast, downregulated gene in *LMNA*^-/-^ cells or LMNA-S22A/S392A-overexpressing cells were not closer to dLASs (**Fig. 2d; Extended Data Fig. 3**). The biased localization of dLASs near genes transcriptionally de-repressed by *LMNA* deletion suggests a link between depolymerized LMNA binding in G2 and transcriptional repression.

**Figure 2.**
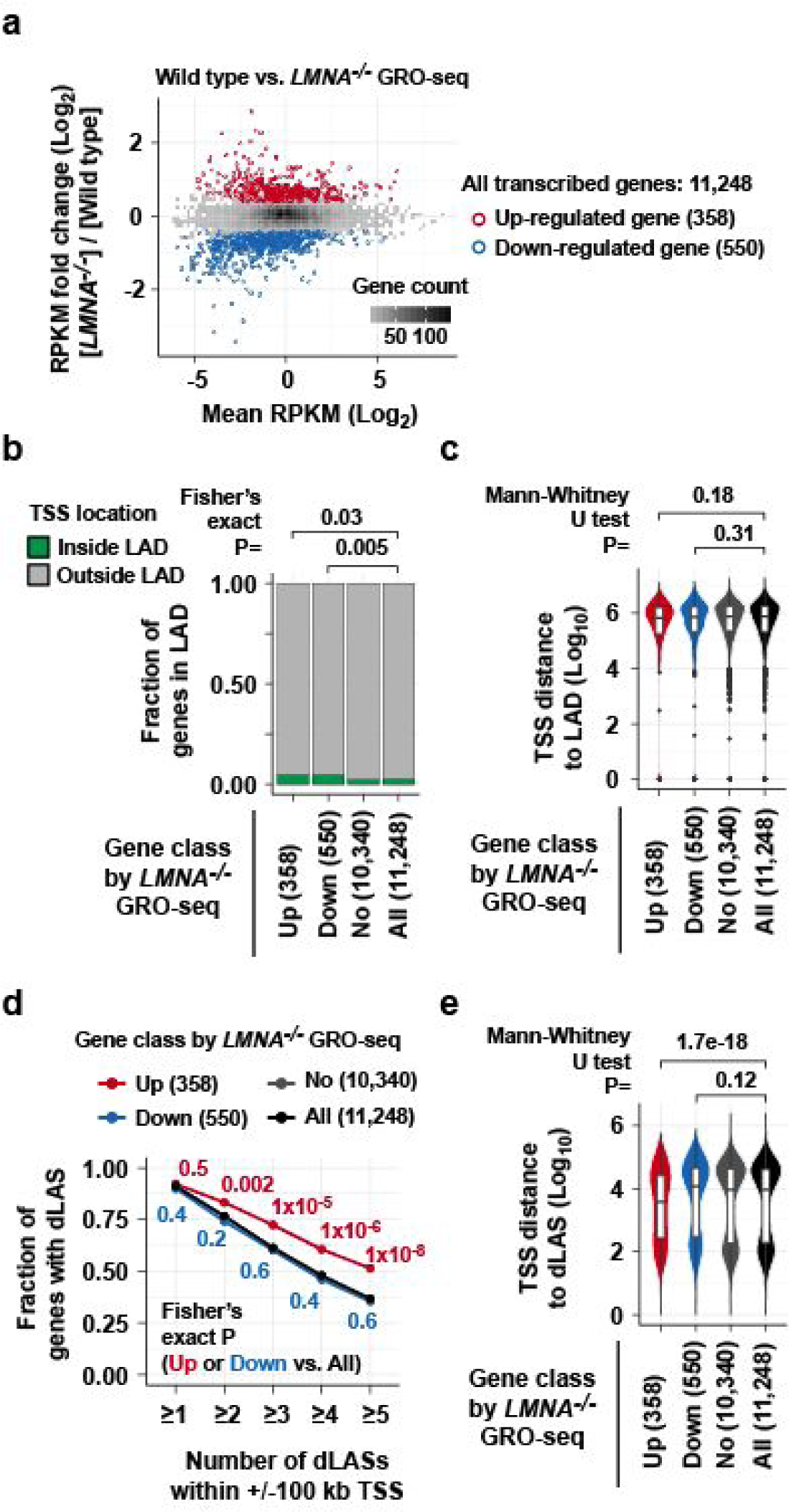
Depolymerized-LMNA-associated sites are over-represented near genes de-repressed by *LMNA* deletion. **a.** MA plot for gene transcription levels (GRO-seq RPKM) in wild-type and *LMNA*^*–/–*^ BJ-5ta cells, highlighting *LMNA*-dependent genes. **b.** Fraction of genes whose TSSs are located inside or outside LADs. Up, *LMNA*^*–/–*^-up genes. Down, *LMNA*^*–/–*^-down genes. No, unchanged genes. All, all transcribed genes. **c.** TSS-to-LAD distance for *LMNA*-dependent genes. **d.** Fraction of genes with dLASs within +/– 100 kb of TSS. Fraction is computed for each dLAS number cutoff indicated at the x-axis. **e.** TSS-to-dLAS distance for *LMNA*-dependent genes.

We hypothesized that depolymerized LMNA contributes to transcriptional repression in G2 in preparation for mitosis. To test this hypothesis, we isolated early G1 (EG1), G1, S, and G2/M cell-cycle stages from asynchronous BJ-5ta cells by FUCCI-based cell sorting^19^ and performed RNA-seq (**Extended Data Fig. 4a–c**). We first confirmed expected expression dynamics of known cell-cycle-stage-dependent genes (**Extended Data Fig. 4d–g**). *LMNA*^*–/–*^-up genes revealed a striking cell-cycle-stage-dependent pattern: their mRNA levels underwent a steep reduction at G2/M (**Fig. 3a**). *LMNA*^*–/–*^-down genes did not demonstrate a coherent cell-cycle-stage-dependent pattern (**Fig. 3a**). As a complementary approach, we employed unsupervised clustering, which extracted 4 gene clusters with distinct cell-cycle expression dynamics from all genes (**Extended Data Fig. 4h**). *LMNA*^*–/–*^-up genes were strongly enriched in Clusters 1 and 4, distinguished by peak expression at G1 or S, with a subsequent steep decline at G2/M (P=4×10^−13^; **Fig. 3b**). Direct computation of mRNA fold change between S and G2/M stages further confirmed the strong downregulation of *LMNA*^*–/–*^-up genes at G2/M (P=2×10^−20^; **Extended Data Fig. 4i**). Thus, the transcriptional targets of depolymerized LMNA were downregulated at G2/M in wild-type cells, supporting our hypothesis that depolymerized LMNA contributes to transcriptional repression at G2.

**Figure 3.**
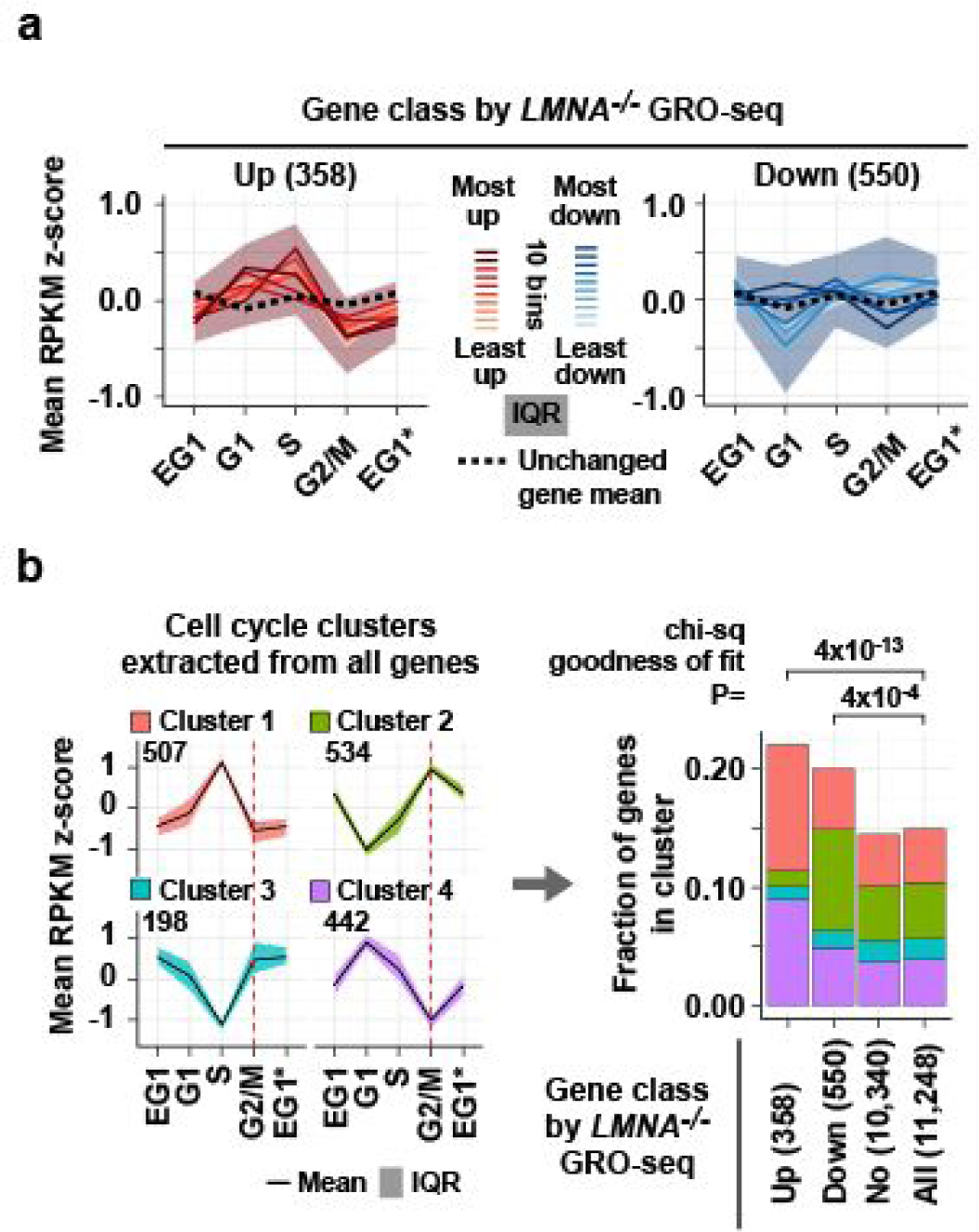
Depolymerized LMNA-target genes are repressed at G2/M. **a.** Expression levels of *LMNA*-dependent genes in wild-type BJ-5ta cells at early G1 (EG1), G1, S, and G2/M cell-cycle stages measured by RNA-seq. *LMNA*-dependent genes are grouped into 10 equally-sized bins based on GRO-seq log_2_ fold change (absolute log_2_([*LMNA*^*–/–*^]/[wild type])) in asynchronous cells. Solid line, within-bin mean RPKM z-score. Shade, interquartile range (IQR) of z-scores for all *LMNA*^*–/–*^-up and *LMNA*^*–/–*^-down genes. Dashed line, mean z-score of *LMNA*^*–/–*^-unchanged genes. *EG1 data is replicated to show G2/M to EG1 change. **b.** (Left) Expression dynamics of the 4 cell-cycle gene clusters extracted from the RNA-seq data (**Extended Data Fig. 4h**). (Right) Fraction of genes assigned to the 4 cell-cycle gene clusters.

We hypothesized two mechanisms by which depolymerized LMNA represses transcription at G2. Depolymerized LMNA may (a) inhibit recruitment of RNA Polymerase II (Pol II) to promoters and/or (b) prevent transcriptional elongation. To assess these possibilities, we examined the GRO-seq signal distribution along genes, which reflects the Pol II abundance^20^. In *LMNA*^*–/–*^-up genes, the difference in GRO-seq signals between wild-type and *LMNA*^*–/–*^ cells was modest at the TSS-proximal region (TSS to +500 bp) (**Fig. 4a**). However, significantly increased GRO-seq signals were present along gene bodies of *LMNA*^*–/–*^ cells compared to the wild-type cells (**Fig. 4a**), suggesting that *LMNA* deletion resulted in excessive Pol II elongation rather than excessive recruitment. The largest increase of GRO-seq signals in *LMNA*^*–/–*^ cells occured within the first 5 kb of TSS, suggesting that increased Pol II pause release contributed to increased transcriptional elongation (**Fig. 4a**). A similar increase of GRO-seq signals along gene bodies was also observed in genes de-repressed by LMNA phospho-deficient mutant overexpression (**Fig. 4b**). Consistent with the presence of dLASs at nearly all transcribed genes, even genes not classified as *LMNA*-dependent demonstrated a significant increase of GRO-seq signals within the first 5 kb of TSS (“No change” in **Fig. 4a**). These data suggest that depolymerized LMNA restricts transcriptional elongation likely by promoting Pol II pausing.

**Figure 4.**
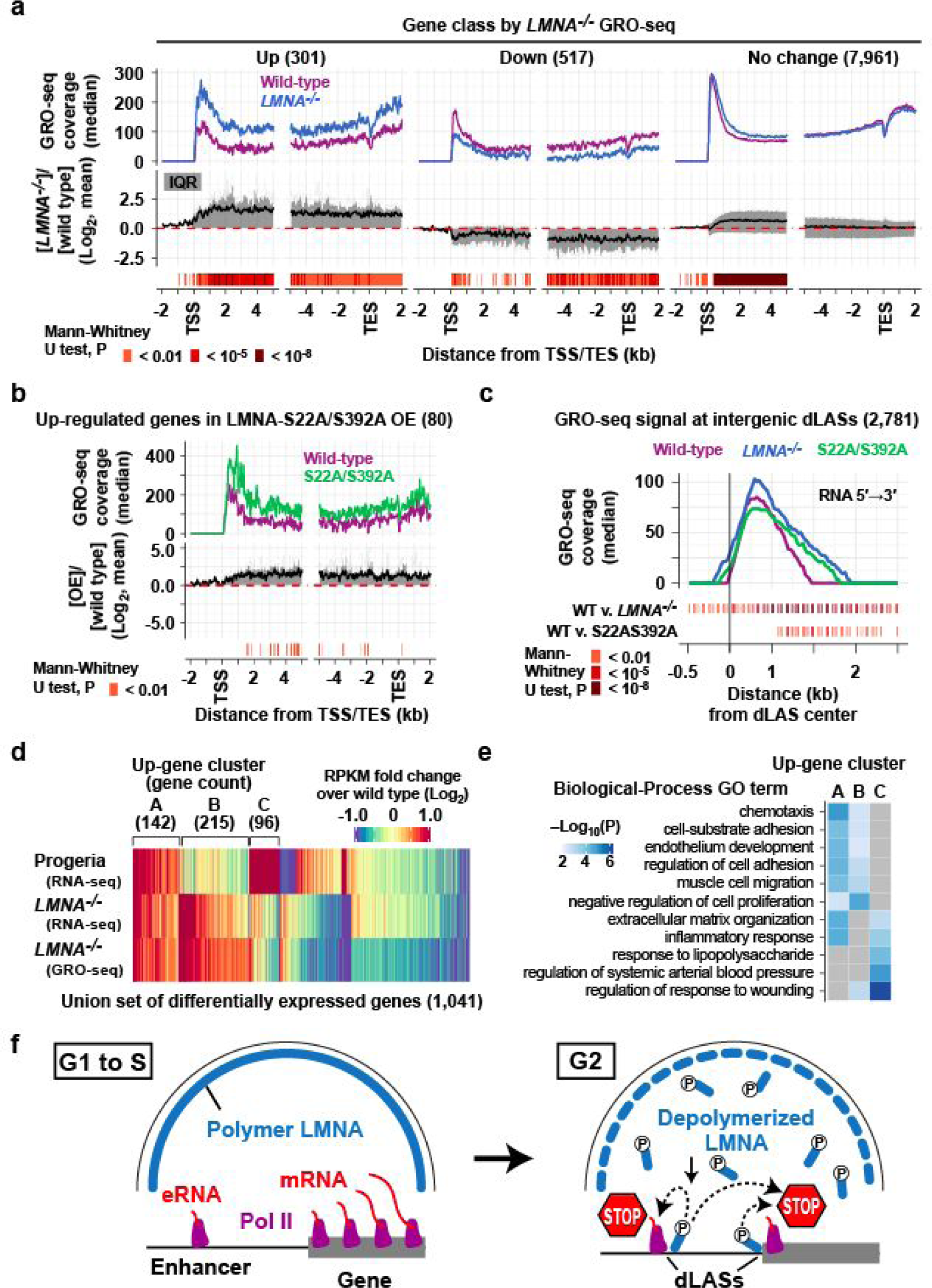
*LMNA* deletion results in excessive transcriptional elongation genome-wide. **a.** (Top) GRO-seq read coverage along genes. Coverage for 5′ to 3′ GRO-seq reads relative to gene orientation are computed in 50-bp windows with a 25-bp offset. Genes with a gene-body size larger than 10 kb are analyzed. (Middle) Distribution of log_2_ signal difference. Line, mean. Shade, IQR. (Bottom) Windows with statistically significant coverage difference. **b.** Same as **a**, but genes up-regulated by over-expression of phospho-deficient LMNA-S22A/S392A are shown. **c.** Same as **a**, but GRO-seq read coverage at intergenic distal (>1 kb TSS) dLASs with minimum 10 read coverage are shown. Coverage is computed for each genomic strand. **d.** RPKM fold change for a union of *LMNA*-dependent genes and genes dysregulated in progeria patient fibroblasts. Capital letters indicate three clusters of up-regulated genes. **e.** Gene ontology (GO) terms enriched in up-regulated gene clusters in **d**. **f.** Model. Depolymerized LMNA targets active regulatory elements to promote Pol II pausing in G2 in preparation for mitotic transcriptional quiescence.

Given that dLAS localizes at putative active enhancers genome-wide, we asked if Pol II-dependent enhancer transcription was affected by *LMNA* deletion. At intergenic, gene-distal dLASs with measurable GRO-seq read coverage (2,781 dLASs), *LMNA*^*–/–*^ and LMNA phospho-deficient mutant overexpression cells showed longer “tails” of GRO-seq signals relative to wild-type cells (**Fig. 4c**), indicating an increase of elongating Pol II. Thus, abnormally increased elongating Pol II is widespread at promoters and enhancers in *LMNA*^*–/–*^ cells and phospho-deficient LMNA mutant overexpressing cells.

In Hutchinson-Gilford progeria, a premature aging disorder caused by a heterozygous splice-site mutation in *LMNA*^*5*^, the mutant LMNA protein is permanently farnesylated^21^, and consequently, remains associated with the nuclear membrane during the mitosis and does not depolymerize^22^. We therefore hypothesized that progeria-patient cells exhibit defects in depolymerized LMNA-mediated transcriptional repression. RNA-seq in primary skin fibroblast cells derived from two progeria patients identified 139 up-regulated and 57 down-regulated genes relative to primary skin fibroblasts from two normal individuals (**Fig. 4d; Extended Data Fig. 2**). These up-regulated genes in progeria cells significantly overlapped *LMNA*^*–/–*^-up genes (29 genes, 21%; P=4×10^−16^) while down-regulated genes in progeria did not significantly overlap *LMNA*^*–/–*^-up or *LMNA*^*–/–*^-down genes (**Fig. 4d; Extended Data Fig. 2**). Genes commonly up-regulated in the progeria-patient and *LMNA*^*–/–*^ fibroblasts (Cluster A) were enriched for gene ontology terms related to cell migration/adhesion, inflammatory response, and extracellular matrix (ECM) organization (**Fig. 4e**), which may be related to severe fibrosis of arterial adventitia in progeria, a feature unique to arteriosclerosis in progeria^23^. Thus, a defect of depolymerized LMNA-mediated transcriptional repression may contribute to gene expression defects in progeria-patient fibroblasts and progeria pathogenesis.

Several mechanisms have been proposed for mitotic transcriptional quiescence, including termination of elongating Pol II^24^, clearance of paused Pol II^25^, inactivation of TFIID^26^, and physical exclusion of transcription factors^27^. These proposed mechanisms operate during mitosis^24–27^. Because LMNA depolymerization occurs in G2 before chromatin condensation, the depolymerized LMNA-mediated mechanism is likely upstream in the sequence of the events that ultimately causes transcriptional quiescence in mitosis. The large number of LMNA molecules stored as polymers in the nuclear lamina during interphase may afford rapid and genome-wide transcriptional pausing through the rapid depolymerization of LMNA during G2. Collectively, our data support a model in which depolymerized LMNA targets active regulatory elements in G2 to promote Pol II pausing, thereby promoting transcriptional repression preceding mitosis (**Fig. 4f**). Our results thereby provide evidence that two fundamental aspects of animal cell division, nuclear envelope breakdown and mitotic transcriptional silencing, are molecularly coupled by depolymerized LMNA. Lamins therefore have gene regulatory functions outside the nuclear lamina in a depolymerized form, which may be relevant to the pathogenesis of nuclear lamin-related disease.

## METHODS

### Statistics

***Permutation test in Figure 1h:*** For 34,489 dLAS summits (1 bp), a random set of 34,489 genomic locations was selected from the blacklisted-region-filtered genome using *shuffle* function in Bedtools (bedtools 2.25.0)^28^ such that the chromosome distribution of the original 34,489 dLASs was maintained. This process was iterated for 1,000 times. For each iteration, the total number of bases overlapped with a given chromatin state was computed. For each chromatin state, the number of iterations in which the base coverage of permutated dLASs exceeded the actual base coverage of the 34,489 dLASs was counted. If this number is 0 (one-side test for over-representation), we assigned empirical P-value of < 0.05. Essentially, the same computation was performed for 2,178 LADs, except that the random sets of LADs maintained the distribution of the original feature sizes.

***Mann-Whitney U test in Figure 1k*:** The test was performed under the null hypothesis that signal distributions of dLAS-overlapping and non-overlapping sites are equal, with the alternative hypothesis that the two distributions are not equal.

***Fisher’s exact test in Figure 2b & Extended Data Figure 3a:*** The tests were performed under the null hypothesis that, among the 11,248 all transcribed genes, there is no association between being differentially expressed and being inside LADs (the odds ratio equals one) with the alternative hypothesis that there is an association (the odds ratio does not equal one).

***Fisher’s exact test in Figure 2d & Extended Data Figure 3c:*** The tests were performed under the null hypothesis that, among the 11,248 all transcribed genes, there is no association between being differentially expressed and having indicated number of dLASs (the odds ratio equals one) with the alternative hypothesis that there is an association (the odds ratio does not equal one).

***Mann-Whitney U test in Figure 2c, e, & Extended Data Figure 3b, d:*** The test was performed under the null hypothesis that distribution of TSS distances to dLAS or LADs in *LMNA*^*–/–*^-up or *LMNA*^*–/–*^-down genes equals that of all transcribed genes, with the alternative hypothesis that the distributions are not equal.

***Chi-square goodness of fit test in Figure 3b:*** The tests were performed under the null hypothesis that the observed frequencies of the 4 clusters are equal to the frequencies in the all transcribed genes, with the alternative hypothesis that the frequencies are not equal.

***Mann-Whitney U test in Figure 4a–c:*** The test was performed under the null hypothesis that signal distributions of wild-type and *LMNA* mutant cells are equal, with the alternative hypothesis that the distributions are not equal.

***GO-term enrichment in Figure 4e***: P-values indicating GO-term enrichment were computed using Metascape^29^, which uses a cumulative hypergeometric statistical test.

***Student’s t-test in Extended Data Figure 5d:*** The test was performed under the null hypothesis that the mean of cell counts in 7 growth experiments in BJ-5ta cells and that of 7 growth experiments in *LMNA*^*–/–*^ cells are equal, with the alternative hypothesis that the means are not equal.

***Fisher’s exact test in Extended Data Figure 2:*** The tests were performed under the null hypothesis that, among the 11,248 transcribed genes, there is no association between being differentially expressed in one condition and another condition (the odds ratio equals one) with the alternative hypothesis that there is an association (the odds ratio does not equal one).

***Mann-Whitney U test in Extended Data Figure 4i:*** The test was performed under the null hypothesis that the distribution of log_2_ fold change of *LMNA*^*–/–*^-up or *LMNA*^*–/–*^-down genes is equal to that of all transcribed genes, with the alternative hypothesis that the distributions are not equal.

### Cell culture

BJ-5ta (ATCC catalog # CRL-4001) is a TERT-immortalized BJ skin fibroblast cell line that retains normal fibroblast cell growth phenotypes and does not exhibit transformed phenotypes^30^. Generation of the BJ-5ta-derived *LMNA*^-/-^ cell line, BJ-5ta over-expressing *LMNA* mutant transgenes, and BJ-5ta expressing FUCCI fluorescence proteins is described below. Primary skin fibroblasts used are GM07492 (source: thigh of 17-year-old male normal individual, Coriell Cell Repository), GM08398 (source: inguinal area of 8-year-old male normal individual, Coriell Cell Repository), AG11498 (source: thigh of 14-year-old male Hutchinson-Gilford progeria patient, Coriell Cell Repository), and HGADFN167 (source: posterior lower trunk of 8-year-old male Hutchinson-Gilford progeria patient, Progeria Research Foundation).

BJ-5ta, the BJ-5ta-derived *LMNA*^-/-^ cell line, and BJ-5ta overexpressing *LMNA* mutant transgenes were cultured in high-glucose DMEM (Gibco, 11965-092) containing 9% fetal bovine serum (FBS), 90 U/mL penicillin, 90 µg/mL streptomycin streptomycin at 37°C under 5% CO_2_. Primary skin fibroblasts were cultured in MEM Alpha (Gibco, 12561-056) containing 9% fetal bovine serum (FBS), 90 U/mL penicillin, 90 µg/mL streptomycin streptomycin at 37°C under 5% CO_2_.

### LMNA –/– cell line

Briefly, wild-type BJ-5ta cells were transduced with two lentivirus vectors, each of which expressed *S. pyogenes* Cas9 and one of two short-guide RNAs (sgRNA1 and sgRNA3), both of which target the exon 1 of the *LMNA* gene. The transduced cells were clonally expanded. A clone (clone ID cc1170-1AD2) that carries *LMNA* nullizygous mutations (**Extended Data Fig. 5a, b**) and lacks Lamin A and Lamin C protein expression (**Extended Data Fig. 5c**) was obtained. While the *LMNA*^*–/–*^ cell line demonstrated less robust proliferation compared with wild-type cells (**Extended Data Fig. 5d**), cell cycle profiles of wild-type BJ-5ta and the *LMNA*^*–/–*^ cell line were indistinguishable (**Extended Data Fig. 5e**).

In detail, we first introduced DNA sequences for sgRNA1 (oligonucleotides KI223 and KI224; **Supplementary Data 1**) or sgRNA3 (oligonucleotides KI227 and KI228; **Supplementary Data 1**) into lentivirus vector LentiCRISPRv2 (a gift from Feng Zhang; Addgene plasmid # 52961)^31^, yielding pKI9 (for sgRNA1) and pKI13 (for sgRNA3). HEK293FT cells were transfected with pKI9 or pKI13 together with packaging vectors psPAX2 (a gift from Didier Trono; Addgene plasmid # 12260) and pCMV-VSV-G (a gift from Bob Weinberg; Addgene plasmid # 8454)^32^ to produce lentivirus. To transduce BJ-5ta cells with the lentivirus, BJ-5ta cell culture was mixed with cleared tissue-culture supernatant for sgRNA1 lentivirus and that for sgRNA3 (each at 0.25 dilution) in the presence of 7.5 µg/mL polybrene. Successfully transduced cells were selected by 3 µg/mL puromycin and seeded to 10-cm dishes with a density of 100 cells per dish. Clonal populations were expanded and analyzed by western blotting for Lamin A and Lamin C protein expression. One clonal line (cc1170-1AD2) lacked Lamin A and Lamin C protein expression (**Extended Data Fig. 5c**). This cell line possesses nullizygous mutations at the sgRNA1-target site (**Extended Data Fig. 5a**).

### LMNA overexpression

Briefly, for stable expression of *LMNA* transgenes, lentivirus vectors containing cDNA for 3xTy1-tagged LMNA or the isoform C of LMNA (LMNC) or LMNA mutants were transduced to wild-type BJ-5ta cells, and transduced cell populations were selected by puromycin. These transgenes were under the control of the EF1 alpha promoter.

In detail, we first cloned cDNAs for 3xTy1-tagged LMNA, its C-isoform (LMNC) and LMNA phosphorylation-site mutants into lentivirus vector pCDH-EF1α-MCS-PGK-GFP-T2A-Puro (CD813A-1, System Biosciences). cDNA sequences for LMNA, LMNC, and phospho mutants were cloned from cDNA expression plasmids described previously ^16^ (gifts from Drs. Robert Goldman and John E Eriksson). The 3xTy1 tag was incorporated using PCR (oligonucleotides KI253, KI243, and KI244; **Supplementary Data 1**). Lentivectors for Ty1-LMNA (pKI36), Ty1-LMNC (pKI28), Ty1-LMNA-S22A/S392A (pKI111), and Ty1-LMNA-S22D/S392D (pKI114) were transfected to HEK293FT cells to produce lentiviruses. Cleared tissue culture supernatant for Ty1-LMNA and Ty1-LMNC viruses were used in 1/2 dilution, and that for Ty1-LMNA-S22A/S392A and Ty1-LMNA-S22D/S392D viruses were used in 1/64 dilution to transduce wild-type BJ-5ta cells. Transduced cell populations were selected under 3 µg/mL puromycin. The cell IDs for obtained cell populations are cc1194-4-1 (Ty1-LMNA OE); cc1194-5-1 (Ty1-LMNC OE); cc1255-2-6 (Ty1-LMNA-S22A/S392A OE); and cc1255-3-6 (Ty1-LMNA-S22D/S392D).

### FUCCI-expressing cells

For stable expression of FUCCI-fluorescent proteins, we used FUCCI4 lentivirus vectors^19^, pLL3.7m-Clover-Geminin^1-110^-IRES-mKO2-Cdt^30-120^(Addgene plasmid #83841) and pLL3.7m-mTurquoise2-SLBP^18-126^-IRES-H1-mMaroon1 (Addgene plasmid #83842) (gifts from Michael Lin). HEK293FT cells were transfected with these lentiviral vectors to produce Clover/mKO2 and mTurquoise/mMaroon1 lentiviruses individually. BJ-5ta cells were co-transduced with the virus supernatant for the two viruses (0.017:1 dilution each). A BJ-5ta cell population positive for both Clover and mMaroon1 fluorescence (BJ-5ta^FUCCI^) was selected using fluorescence-activated cell sorting (FACS) and maintained (cell population ID cc1357-2ss).

### FUCCI cell sorting

In the FUCCI4 system, G0/G1-phase cells are positive for mKO2-Cdt^30-120^; S-phase cells are positive for mKO2-Cdt^30-120^, mTurquoise2-SLBP^18-126^, and Clover-Geminin^1-110^; G2/M cells are positive for Clover-Geminin^1-110^; early G1 cells are negative for all of these three marker proteins^19^ (**Extended Data Fig. 4a**). Before sorting, asynchronously cultured BJ-5ta^FUCCI^ cells were cultured at a 50-70% density to prevent G0 entry. For FACS, cells were trypsinized. FACS was performed on BD FACS Aria II (biological replicate IDs 2834 and 2847) or BD FACS Aria IIIu (biological replicate ID 2858) (**Extended Data Fig. 4b**). With FACS, mKO2-positive/Clover-negative population was sorted as G1-phase cells, mKO2-negative/Clover-positive population was sorted as G2/M-phase cells, mKO2-negative/Clover-negative population was sorted as EG1-phase cells, and mKO2-positive/Clover-positive/mTurquoise2-positive population was sorted as S-phase cells (**Extended Data Fig. 4b**). Pre-sorting and post-sorting cells were maintained in the culture medium on ice. Sorted cells were pelleted and resuspended in Trizol LS (Invitrogen 10296028) for RNA-seq. We performed three biological replicates of BJ-5ta^FUCCI^ cell culture, each of which generated 4 RNA-seq data sets corresponding to the 4 cell-cycle stages (EG1, G1, S, and G2/M).

### Immunofluorescence

Cells were grown on uncoated coverslips under the standard culture condition (see *Cell Culture*) and fixed in PHEM buffer (60 mM PIPES-KOH pH7.5, 25 mM HEPES-KOH pH7.5, 10 mM EGTA, 4 mM MgSO_4_) containing 4% formaldehyde and 0.5% Triton for 10 min at 37°C. Cells on coverslips were blocked in PBS with 1% slim milk and 5% goat serum and incubated with primary antibodies. Antibodies used in immunofluorescence are: rabbit monoclonal anti-phospho-Ser22-LMNA antibody D2B2E (Cell Signaling 13448S, Lot # 1); mouse monoclonal anti-unphospho-Ser22-LMNA antibody E1 (Santa Cruz Biotechnology sc-376248, Lot # H2812); and mouse monoclonal anti-phospho-Ser10 histone H3 antibody MA5-15220 (Thermo Fisher MA5-15220, Lot # RH2257952). Cells were incubated with secondary antibodies, counterstained by DAPI, and cured in ProLong Gold mounting medium (Molecular Probes, P36930). Slides were imaged using Leica SP8 confocal microscope with a 63x objective.

### Western blot

Ten micrograms of total proteins were separated by SDS-PAGE and transferred to a PVDF membrane. LMNA proteins were detected by rabbit polyclonal anti-LMNA antibody H-110 (Santa Cruz Biotechnology sc-20681, Lot # D1415, 1:5000 dilution). The gel after protein transfer was counter-stained by coomassie to evaluate the loaded protein amount.

### ChIP-seq

Cells in culture dishes were crosslinked in 1% formaldehyde for 15 min, and the reaction was quenched by 125 mM glycine. Cross-linked cells were washed with LB1 (50 mM HEPES-KOH pH7.5, 140 mM NaCl, 1 mM EDTA, 10% glycerol, 0.5% NP40, 0.25% Triton X-100) and then with LB2 (200 mM NaCl, 1 mM EDTA, 0.5 mM EGTA, and 10 mM Tris-HCl pH 8.0). For cell-cycle-stage specific ChIP, cross-linked cells were stained by DAPI (5 µg/mL) in LB1 for 5 min at RT, sorted based on the cellular DNA content using BD FACS Aria II or BD FACS Aria IIIu, and then washed with LB2. Chromatin was extracted by sonication in LB3-Triton (1 mM EDTA, 0.5 mM EGTA, 10 mM Tris-HCl pH 8, 100 mM NaCl, 0.1% Na-Deoxycholate, 0.5% N-lauroyl sarcosine, 1% Triton). LB1, LB2, and LB3-Triton used during wash and extraction were supplemented with 1x protease inhibitor cocktail (Calbiochem 539131) and 100 nM phosphatase inhibitor Nodularin (Enzo ALX-350-061). The cell extract was cleared by 14,000 g centrifugation for 10 min. To evaluate chromatin extraction efficiency, DNA were extracted from aliquots of the supernatant and pellet by phenol chloroform isolation. We ensured that ≥95% of genomic DNA is solubilized in the supernatant. An aliquot of cell extract was saved for input DNA sequencing.

For ChIP, cell extract from one million cells was incubated with antibodies in a 200-µL reaction for ≥12 hours. Antibodies used in ChIP are: rabbit monoclonal anti-phospho-Ser22-LMNA antibody D2B2E (Cell Signaling 13448S, Lot # 1; 5 µL per IP); mouse monoclonal anti-unphospho-Ser22-LMNA antibody E1 (Santa Cruz Biotechnology sc-376248, Lot # H2812; 10 µL per IP); mouse monoclonal anti-acetyl-Lys27 histone H3 antibody (Wako MABI0309, Lot # 14007; 2 µL per IP); mouse monoclonal anti-trimethyl-Lys4 histone H3 antibody (Wako MABI14004, Lot # 14004; 2 µL per IP); mouse monoclonal anti-Ty1 antibody (Diagenode C15200054, Lot # 005; 1 µL per IP). Immunocomplex was captured by Protein A-conjugated sepharose beads (for rabbit antibodies) or Protein G-conjugated magnetic beads (for mouse antibodies) and washed. Immunoprecipitated DNA was reverse-crosslinked and used to construct high-throughput sequencing libraries using NEBNext Ultra DNA Library Prep Kit (New England Biolabs, E7370). DNA libraries were processed on a Illumina HiSeq machine for single-end sequencing. Data processing steps are described in *ChIP-seq data processing*.

### ATAC-seq

One hundred thousand trypsinized cells were incubated with ATAC hypotonic buffer (10 mM Tris pH 7.5, 10 mM NaCl, 3 mM MgCl_2_) at 4°C for 10 min during 500 g centrifugation. Cells were incubated in Tagmentation mix (Tagmentation DNA buffer Illumina 15027866; Tagmentation DNA enzyme Illumina 15027865) at 37°C for 30 min. Purified DNA was used to construct high-throughput sequencing libraries using NEBNext High-Fidelity 2x PCR Master Mix (New England Biolabs M0541). DNA libraries were processed on a Illumina NextSeq machine for paired-end 41-nt sequencing. Data processing steps are described in *ATAC-seq data processing*.

### GRO-seq

Nuclei isolated from trypsinized cells are resuspended in Nuclear Storage buffer (50 mM Tris pH 8.0, 0.1 mM EDTA, 5 mM MgCl_2_, 40% glycerol, RNase inhibitor). An equal volume of 2x NRO buffer (10 mM Tris pH 8.0, 5 mM MgCl_2_, 1 mM DTT, 300 mM KCl, 0.5 mM ATP, 0.5 mM GTP, 0.5 mM BrUTP, 2 µM CTP) was added to the nuclear suspension and incubated for 8 min at 30°C for nuclear run-on reaction. During the incubation, sarkosyl was added to the reaction (0.5% final concentration) 4 min after the initiation of the reaction. RNAs were purified from the reaction by Trizol LS (Invitrogen 10296028) followed by isopropanol precipitation. RNAs were treated with TurboDNase (Ambion AM18907) and fragmented by Fragmentation Buffer (Ambion AM8740). BrU-incorporated RNA fragments were immunoprecipitated with mouse monoclonal anti-BrdU antibody 3D4 (BD Biosciences 555627) and used to construct DNA sequencing libraries using NEBNext Ultra II Directional RNA Library Prep kit (New England Biolabs E7760). DNA libraries were processed on a Illumina HiSeq machine for single-end 50-nt sequencing. Data processing steps are described in *GRO-seq data processing*.

### RNA-seq

For non-FUCCI cells, total RNAs were purified by Trizol (Invitrogen 15596026). After DNase treatment, mRNAs were isolated using NEBNext Poly(A) mRNA Magnetic Isolation Module (New England Biolabs E7490) and fragmented by Fragmentation Buffer (Ambion AM8740). cDNAs were synthesized using SuperScript II (Invitrogen 18064014) and used to prepare non-directional high-throughput sequencing libraries using NEBNext Ultra DNA Library Prep Kit (New England Biolabs, E7370). For FUCCI cells, total RNAs were purified by Trizol LS (Invitrogen 10296028). After DNase treatment, mRNAs were isolated using NEBNext Poly(A) mRNA Magnetic Isolation Module (New England Biolabs E7490). Fragmentation, cDNA synthesis, and directional library construction were performed using NEBNext Ultra II Directional RNA Library Prep kit (New England Biolabs E7760). The both types of libraries were processed on the Illumina HiSeq platform for single-end 50-nt sequencing. Data processing steps are described in *RNA-seq data processing*.

### CNV detection

An initial visual inspection of ChIP-seq signals in a genome browser led us to suspect a copy number variation (CNV) at a large genomic region in the right end of the chromosome I, which exhibited overall weaker ChIP and input signals in the *LMNA*^*–/–*^ cell line compared with the wild-type BJ-5ta cell line. To systematically identify potential CNVs in the *LMNA*^*–/–*^ cell line, we used CNV-seq^33^, a computational pipeline to detect CNVs from high-throughput sequencing data. CNV-seq was run using input sequencing data for wild-type BJ-5ta (ID KI481) and *LMNA*^*–/–*^ cells (ID KI489) with the following set of parameters: [–genome human –global-normalization –log2-threshold 0.5 –minimum-windows-required 3]. CNV-seq identified 3,717 candidate CNV windows (window size, 25,783 bp; step size, 12,893 bp). To reduce false positive windows due to low input sequencing coverage, a positive window was required to contain the sum of aligned reads of KI481 and KI489 greater than 200, resulting in 3,073 candidate windows. Overlapping windows or neighboring windows spaced within 500 kb were merged, and those smaller than 500 kb in size were removed. This yielded 5 candidate CNVs (24 Mb in chr1; 15 Mb in chr 4; 2.7 Mb in chr19; 683 kb in chr2; and 528 kb in chrX), which included the suspected region on chr1. These 5 CNVs were added to the list of blacklisted regions (see *Blacklisted region* section) and excluded from all analyses.

### Gene annotation

The Gencode V19 “Basic” gene annotation was downloaded from the UCSC genome browser. Of the total 99,901 transcripts in the list, we retained transcripts that met all of the following requirements: 1) “gene type” equals “transcript type”; 2) “transcript type” is either *protein_coding* or *antisense* or *lincRNA*; and 3) “transcript ID” appears only once in the list. This processing yielded 75,968 transcripts. To select one transcribed unit per gene locus, the 75,968 transcripts were grouped by “gene symbol,” and within the group, transcripts were sorted by the “exon count” (largest first), then by transcribed region “length” (largest first), then by the alphanumeric order of the “transcript ID” (smallest first), and the transcript that appeared first in the group was chosen to represent the transcribed unit of that gene. In this processing, in general, a gene is represented by a transcript with a largest number of annotated exons or largest in size among other associated transcripts. After removing genes located within the blacklisted regions, we obtained 31,561 genes, which included 19,469 “protein_coding” genes.

### ChIP-seq data processing

ChIP-seq experiments and sequencing depth are listed in **Supplementary Table 1**. ChIP-seq reads were mapped to the hg19 human reference genome using Bowtie2 with the default “--sensitive” parameter. Reads with MAPQ score greater than 20 were used in downstream analyses. Reads from biological replicates of ChIP and the corresponding input were processed by MACS2^34^. In MACS2, duplicate reads were removed, and then significantly enriched peak regions were called, and input-normalized perbase coverage of 200 nt-extended reads (fold enrichment data) was computed. Peaks overlapping blacklisted regions (see *Blacklisted regions* section) were removed. MACS2 identified 34,489 enriched sites for pS22-LMNA ChIP-seq in BJ-5ta (P-value cutoff of 10^−20^) (**Supplementary Data 2**); 79,799 sites for H3K27ac ChIP-seq in BJ-5ta (P-value cutoff of 10^−5^) (**Supplementary Data 5**); 18,100 sites for H3K4me3 ChIP-seq in BJ-5ta (P-value cutoff of 10^−5^) (**Supplementary Data 6**); 31,275 sites for Ty1-LMNA ChIP-seq (P-value cutoff of 10^−20^) (**Supplementary Data 7**); 22,411 sites for Ty1-LMNC ChIP-seq (P-value cutoff of 10^−20^) (**Supplementary Data 8**); 6,044 sites for Ty1-LMNA-S22A/S392A ChIP-seq (P-value cutoff of 10^−20^) (**Supplementary Data 9**); and 31,392 sites for Ty1-LMNA-S22D/S392D ChIP-seq (P-value cutoff of 10^−20^) (**Supplementary Data 10**).

### ATAC-seq data processing

ATAC-seq experiments and sequencing depth are listed in **Supplementary Table 1**. For alignment, the first 38 nt from the 5′ ends out of the 41-nt reads were used. The rationale of this trimming is that the minimum size of DNA fragments that can be flanked by Tn5 transposition events has been estimated to be 38 bp^35,36^, and therefore, a 41-nt read could contain a part of read-through adaptors. We aligned 38-nt reads to the hg19 reference genome using bowtie2 with following parameters: [-X 2000 --no-mixed --no-discordant --trim3 3]. The center of the active Tn5 dimer is located +4-5 bases offset from the 5′-end of the transposition sites^35,36^. To place the Tn5 loading center at the center of aligned reads, the 5′-end of the plus-strand and minus-strand reads were shifted 4 nt and 5 nt in the 5′-to-3′ direction, respectively, and the shifted end (1 nt) was extended +/–100 nt. To generate background datasets that capture local bias of read coverage, the shifted read ends were extended +/–5,000 nt and used to construct local lambda background file. This local lambda file and the Tn5 density file were processed by MACS2’s function “bdgcomp” to generate the fold-enrichment file for Tn5 density and by function “bdgpeakcall” to identify regions with statistically significant Tn5 enrichment (ATAC peaks). At P-value cutoff of 10^−10^, we obtained 73,933 ATAC peaks (**Supplementary Data 4**).

### RNA-seq data processing

RNA-seq experiments and sequencing depth are listed in **Supplementary Table 1**. RNA-seq reads were aligned to the hg19 human reference genome using Tophat2 with the default parameter set. Reads with MAPQ score greater than 50 were used in downstream analyses. To compute RPKM (Reads Per Kilobase of transcript per Million mapped reads) for each transcribed region (see *Gene annotation* section), the sum of perbase read coverage in exons was computed, and this was divided by the read length (50), then by the sum of the exon size in kb, and then by the total number of reads in million. In directional RNA-seq datasets, only reads whose direction matches that of the gene model were used to compute RPKM. To identify differentially expressed genes, we used DESeq2 (version 1.14.1) with unnormalized read coverage for the 31,561 transcribed regions (see *Gene annotation* section). We also used DESeq2 to compute log_2_ fold change for comparisons indicated in **Extended Data Fig. 2**. We called differentially expressed genes under the adjusted P-value cutoff of 0.05 with default parameters. The number of differentially expressed genes is listed in **Extended Data Fig. 2**. The differentially expressed genes are listed in **Supplementary Data 11**.

### GRO-seq data processing

GRO-seq experiments and sequencing depth are listed in **Supplementary Table 1**. GRO-seq reads were mapped to the hg19 human reference genome using Tophat2 with the default parameter set. Reads with MAPQ score greater than 50 were used in downstream analyses. RPKM scores were computed as in directional RNA-seq above except that the sum of perbase read coverage was computed for the entire transcribed regions (instead of exons) and that the size of the transcribed region (instead of the sum of exon lengths) was used for normalization. To obtain replicate-combined depth-normalized read coverage files (bedgraph) for each genomic strand, the perbase coverage was computed from the replicated-combined reads and divided by the total number of reads from all replicates.

### LADs

LADs were defined using UnP-LMNA ChIP-seq data in BJ-5ta. The hg19 genome was segmented into 5-kb non-overlapping windows, and for each window, the sum of perbase read coverage (from replicate-combined reads) was computed for UnP-LMNA ChIP-seq in BJ-5ta and the corresponding input. The coverage was normalized by sequencing depth. The depth-normalized coverage was used to compute per-window log_2_ ratios of ChIP over input. We then created 100 kb windows with the 5-kb step size genome-wide. For each 100-kb window, if every one of the 20 constituting 5-kb windows had a positive log_2_ ratio and the mean log_2_ ratios of the constituting 5-kb windows was greater than 0.5, this 100-kb window was further processed. The qualified 100-kb windows were merged if overlapping. After filtering regions overlapping blacklisted regions (see *Blacklisted regions* section), we obtained 2,178 regions which we defined as LADs in BJ-5ta (**Supplementary Data 3**).

### Blacklisted regions

Before performing post-alignment analyses, we excluded genes and features (such as ChIP-seq peaks and ATAC-seq peaks) located in certain genomic regions that may cause misinterpretation due to high sequence redundancy, uncertain chromosomal locations, high signal background, haplotypes, or CNVs. The collection of such genomic regions were constructed from the following datasets (see *Public datasets* section): assembly gaps in the hg19 reference genome, ENCODE-defined hg19 blacklisted regions, mitochondrion sequence (chrM), haplotype chromosomes (chr*_*_hap*), unplaced contigs (chrUn_*), unlocalized contigs (chr*_*_random), and CNVs in the *LMNA*^*–/–*^ cell line (see *CNV detection* section). The blacklisted regions are listed in **Supplementary Data 12**.

### Aggregate plot and heatmap

To generate aggregate plots and heatmaps for features, two data files were first generated: a) a window file, which consists of, for each genomic feature, an array of fixed-size genomic windows that cover genomic intervals around the feature; and b) a genome-wide signal file (in bedgraph format). For each genomic window in the window file, all signals within that window were obtained from the signal file, and either mean or max of the signals or sum of the perbase signal (“area”) were computed.

For the signal visualization around dLASs (**Fig. 1f, i; Extended Data Fig. 1b, e, g**), an array of 250-bp windows with a 50-bp offset that covered a 10-kb region centered around each of 34,489 dLAS summits or 73,933 ATAC site summits was generated, and the mean of within-window signals was computed from replicate-combined input/background-normalized fold enrichment files.

For the heatmap of LADs (**Extended Data Fig. 1a**), an array of 5-kb windows (without an offset) that covered a LAD body and 250 kb downstream was generated for 1,042 of 2,178 LADs that were extendable to 250 kb downstream without overlapping neighboring LADs. For each window, the mean of within-window signals was computed from replicate-combined input-normalized fold enrichment files.

For the aggregate plots of GRO-seq signals at TSSs (**Fig. 4a, b**), an array of 50-bp windows with 25-bp offset that covered –2 kb to +5 kb of TSS was generated for every gene, and the within-window sum of perbase read coverage for the same-direction reads was computed from replicate-combined depth-normalized strand-specific perbase read coverage file. Only genes whose gene-body size was greater than 10 kb were visualized.

For the aggregate plot of GRO-seq signals at dLASs (**Fig. 4c**), dLASs located in distal intergenic regions (≥1kb TSS excluding gene body) and with the sum of perbase read coverage greater than 500 were used for analysis. For each of the qualified dLASs, the same size windows as the TSS analysis that covered –2 kb to +5 kb of dLAS was generated for every dLAS for each genomic orientation, and the within-window sum of perbase read coverage was computed as in the TSS analysis.

### FUCCI RNA-seq normalization

We performed three biological replicates of BJ-5ta^FUCCI^ cell culture, each of which generated 4 RNA-seq data sets corresponding to the 4 cell-cycle stages (EG1, G1, S, and G2/M). For each gene for each biological replicate, the 4 RPKM scores corresponding to the 4 cell-cycle stages were z-transformed (z=[RPKM-*m*]/*sd*, where *m* and *sd* are the mean and standard deviation of the 4 RPKM scores, respectively). The z-transformed RPKM (termed “RPKM z-score”; **Fig. 3a, b; Extended Data Fig. 4d–h**) recapitulated the cell cycle dynamics of known cell-cycle dependent genes (**Extended Data Fig. 4d–g**).

### Clustering FUCCI RNA-seq data

To cluster genes by cell cycle dynamics (**Fig. 3b; Extended Data Fig. 4h**), we first selected genes whose expression was variable between cell cycle stages but not variable between biological replicates. For this, we computed, for each gene, coefficient of variation (CV=sd/m, where *m* is the mean and *sd* is the standard deviation of RPKM scores) between biological replicates for each cell-cycle stage (CV^stage^), and CV across all 12 samples (CV^all^). We then identified, for each gene, the largest between-replicate CV (CV^stage.max^) and computed log_2_ [CV^all^]/[CV^stage.max^]. This ratio reflects expression variability among cell cycle stages while taking between-replicate variability into account. Across genes, we z-transformed this ratio (“variability index”), and genes with the variability index greater than 1 and the minimum RPKM score across the 12 samples greater than 0 were considered “variable” between cell cycle stages and used in the following K-means clustering (2,020 of 31,561 genes). K-means clustering was performed on RPKM z-scores of these 2,020 genes using the *kmeans* function in R with set.seed(109) and the following parameters: [centers=4, iter.max=50, nstart=50]. Of the 2,020 genes, we used a set of 1,681 genes which were protein-coding genes and among “all transcribed” class of genes (**Fig. 3b** and **Extended Data Fig. 4h**). The clustering assignment is indicated in **Supplementary Data 11**.

### Clustering up-regulated genes

The union (1,041 genes) of *LMNA*-dependent genes (908 genes) and differentially expressed genes in progeria-patient cells (196 genes) was used in the clustering (**Extended Data Fig. 2**). For the 1,041 genes, we constructed a data matrix consisting of log_2_ fold-change scores for [GRO-seq *LMNA*^*–/–*^]/[GRO-seq wild-type], [RNA-seq *LMNA*^*–/–*^]/[RNA-seq wild-type], and [RNA-seq progeria]/[RNA-seq normal]. This data matrix was used as the input for K-means clustering with k=10 using the *kmeans* function in R with set.seed(109) and the following parameters: [centers=10, iter.max=50, nstart=50]. Up-regulated genes appeared in Clusters 4, 6, 7, and 9. Cluster 7 (142 genes) appeared up-regulated in both *LMNA*^*–/–*^ and progeria cells (re-annotated as *Cluster A* in **Fig. 4d, e**). Clusters 6 and 9 (215 genes total) appeared up-regulated only in *LMNA*^*–/–*^ cells (re-annotated as *Cluster B* in **Fig. 4d, e**). Cluster 4 (96 genes) appeared up-regulated only in progeria cells (re-annotated as *Cluster C* in **Fig. 4d, e**). The clustering annotation is indicated in **Supplementary Data 11**.

### Gene ontology analysis

Gene ontology (GO) analyses were performed using Metascape^29^. The input data sets were three lists of Gencode 19 Gene Symbols (see *Gene annotation*) for up-regulated gene clusters A, B, or C (see *Clustering up-regulated genes*). These gene lists were analyzed in Metascape using the “Multiple Gene List” mode. For background, we used all transcribed genes (11,248) defined by GRO-seq (see Main text). Biological processes GO terms with minimum gene count 5, P <0.01, enrichment over background >2 were shown in heatmap (**Fig. 4e**).

### Public datasets

Assembly gaps

http://hgdownload.soe.ucsc.edu/goldenPath/hg19/database/gap.txt.gz

Blacklisted Regions

http://hgdownload.cse.ucsc.edu/goldenPath/hg19/encodeDCC/wgEncodeMapability/wgEncodeDacMapabilityConsensusExcludable.bed.gz

Chromatin annotation

http://egg2.wustl.edu/roadmap/data/byFileType/chromhmmSegmentations/ChmmModels/coreMarks/jointModel/final/E126_15_coreMarks_stateno.bed.gz

Genecode Release 19 Gene Model

http://hgdownload.cse.ucsc.edu/goldenPath/hg19/database/wgEncodeGencodeBasicV19.txt.gz

### Dataset availability

High-throughput sequencing datasets are listed in **Supplementary Table 1** and available at Gene Expression Omnibus (http://www.ncbi.nlm.nih.gov/geo/) under the accession number GSE113354.

### Code availability

All of the scripts used in this study will be deposited to a public github website before publication.

## ACKNOWLEDGEMENTS

We thank technical assistance from the University of Chicago Functional Genomics Core, the University of Chicago Cytometry and Antibody Technology Core, the University of Chicago Light Microscopy Core, and Princeton University Genomics Core Facility. K.I. and I.P.M. are funded by NIH R21/R33 1R21AG054770-01A1. K.I. and J.D.L were funded by Progeria Research Foundation..

## AUTHOR CONTRIBUTIONS

K.I. and J.D.L conceived the study. K.I. and S.S. performed the experiments. K.I. analyzed the data and wrote the manuscript. J.D.L. and I.P.M. provided intellectual contributions and participated in manuscript writing.

## MATERIALS AND CORRESPONDENCE

Correspondence should be addressed to Kohta Ikegami at

## SUPPLEMENTARY DATA

### Supplementary Data 1: Oligonucleotide sequences (fasta)

***Supplementary Data 2: Depolymerized LMNA-associated sites (dLASs) (UCSC narrowPeak)***

***Supplementary Data 3: Lamina-associated domains (LADs) (UCSC Bed)***

***Supplementary Data 4: ATAC-seq peaks in BJ-5ta (UCSC narrowPeak)***

***Supplementary Data 5: H3K27ac ChIP-seq peaks in BJ-5ta (UCSC narrowPeak)***

***Supplementary Data 6: H3K4me3 ChIP-seq peaks in BJ-5ta (UCSC narrowPeak)***

***Supplementary Data 7: Ty1-LMNA ChIP-seq peaks (UCSC narrowPeak)***

***Supplementary Data 8: Ty1-LMNC ChIP-seq peaks (UCSC narrowPeak)***

***Supplementary Data 9: Ty1-LMNA-S22A/S392A ChIP-seq peaks (UCSC narrowPeak)***

***Supplementary Data 10: Ty1-LMNA-S22D/S392D ChIP-seq peaks (UCSC narrowPeak)***

***Supplementary Data 11: Differentially-expressed genes and clusters (text)***

***Supplementary Data 12: Blacklisted regions (UCSC Bed)***

## SUPPLEMENTARY TABLE

***Supplementary Table 1: High-throughput sequencing data sets (Microsoft excel)***

---

**Extended Data Figure 1.**
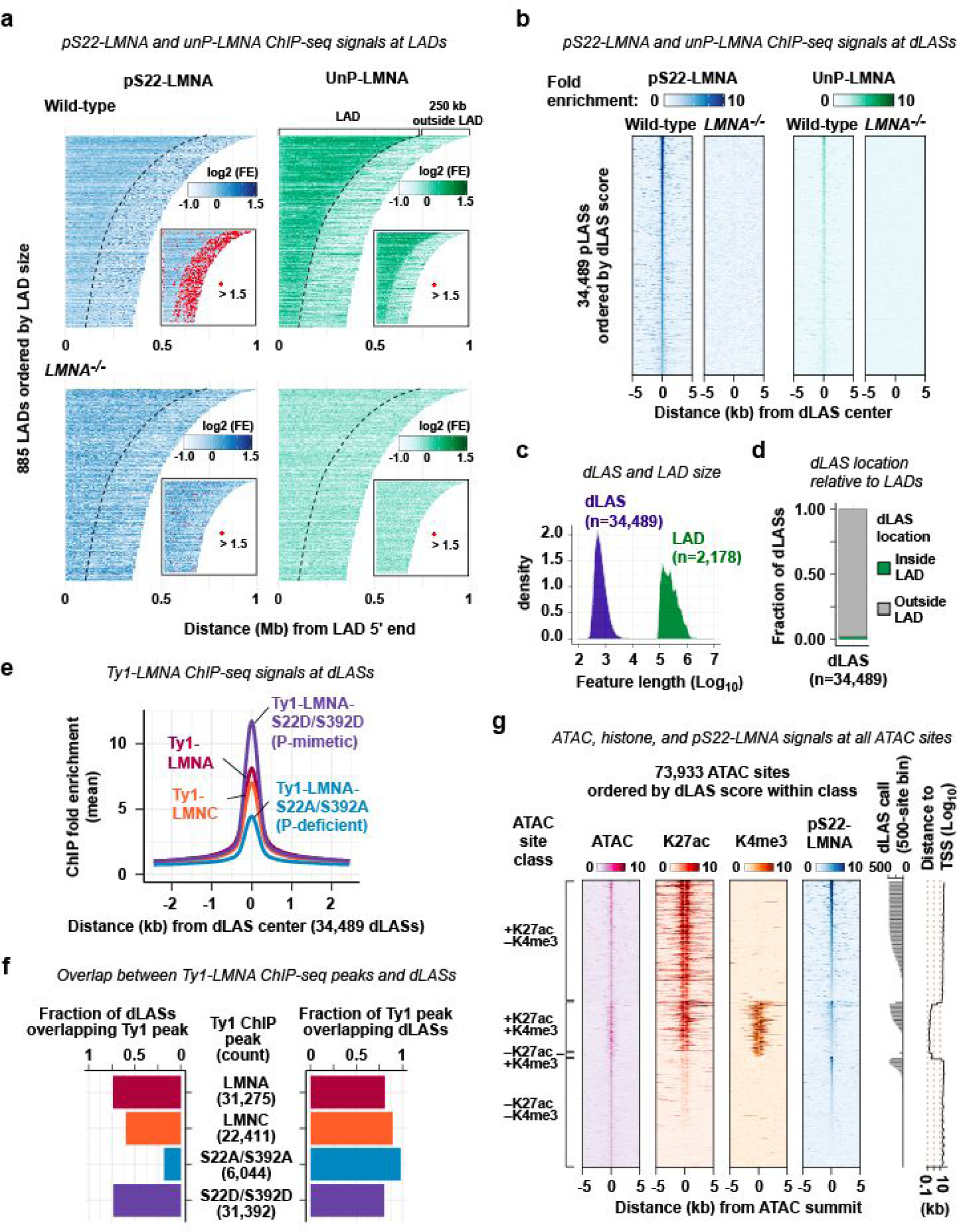
Chracterization of depolymerized LMNA-associated sites. **a.** pS22-LMNA and UnP-LMNA ChIP-seq signals at LADs. LADs are defined by UnP-LMNA ChIP-seq (**Methods**). A subset of LADs (885 of total 2,178) that do not have adjacent LADs within 250 kb downstream of LADs are shown. Inset shows location of strong ChIP-seq signals (marked in red). FE, fold enrichment. **b.** pS22-LMNA and UnP-LMNA ChIP-seq signals at 34,489 dLASs. **c.** Feature size of dLASs and LADs. **d.** Location of dLASs relative to LADs. **e.** Ty1 ChIP-seq signals at dLASs. Ty1 ChIP-seq was performed in BJ-5ta cells overexpressing Ty1-LMNA, Ty1-LMNC (Lamin C), phospho-deficient Ty1-LMNA-S22A/S392, or phospho-mimetic Ty1-LMNA-S22D/S392D. **f.** (Left) Fraction of dLASs overlapping anti-Ty1 ChIP-seq peaks in the overexpression cells shown in **e**. (Right) Fraction of anti-Ty1 ChIP-seq peaks overlapping dLASs. **g.** ATAC-seq, K27ac ChIP-seq, K4me3 ChIP-seq, and pS22-LMNA ChIP-seq signals at 73,933 ATAC-defined accessible sites in wild-type BJ-5ta cells.

**Extended Data Figure 2.**
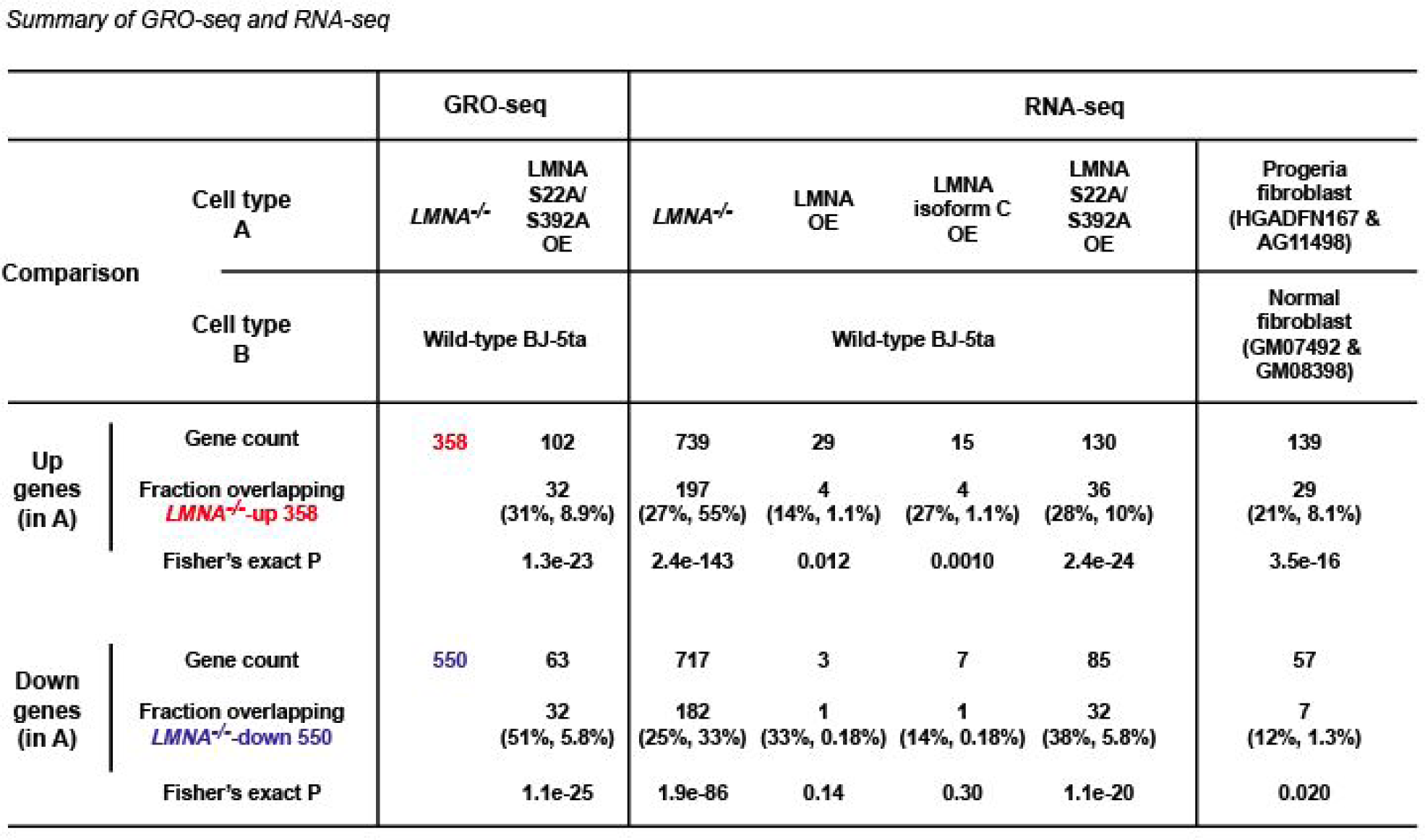
Differentially expressed genes identified in this study. Differentially expressed genes are identified using DESeq2 under adjusted P-value <0.05. Fraction indicates a subset of differentially expressed genes that are also identified as *LMNA*^*–/–*^-up or *LMNA*^*–/–*^-down genes. The first percentage in parentheses indicates the fraction of differentially expressed genes that are identified as *LMNA*^*–/–*^-up or *LMNA*^*–/–*^-down genes. The second percentage indicates the fraction of *LMNA*^*–/–*^-up or *LMNA*^*–/–*^-down genes that are identified as differentially expressed in the indicated comparison. OE, overexpression. Differentially expressed genes are listed in **Supplementary Data 11**.

**Extended Data Figure 3.**
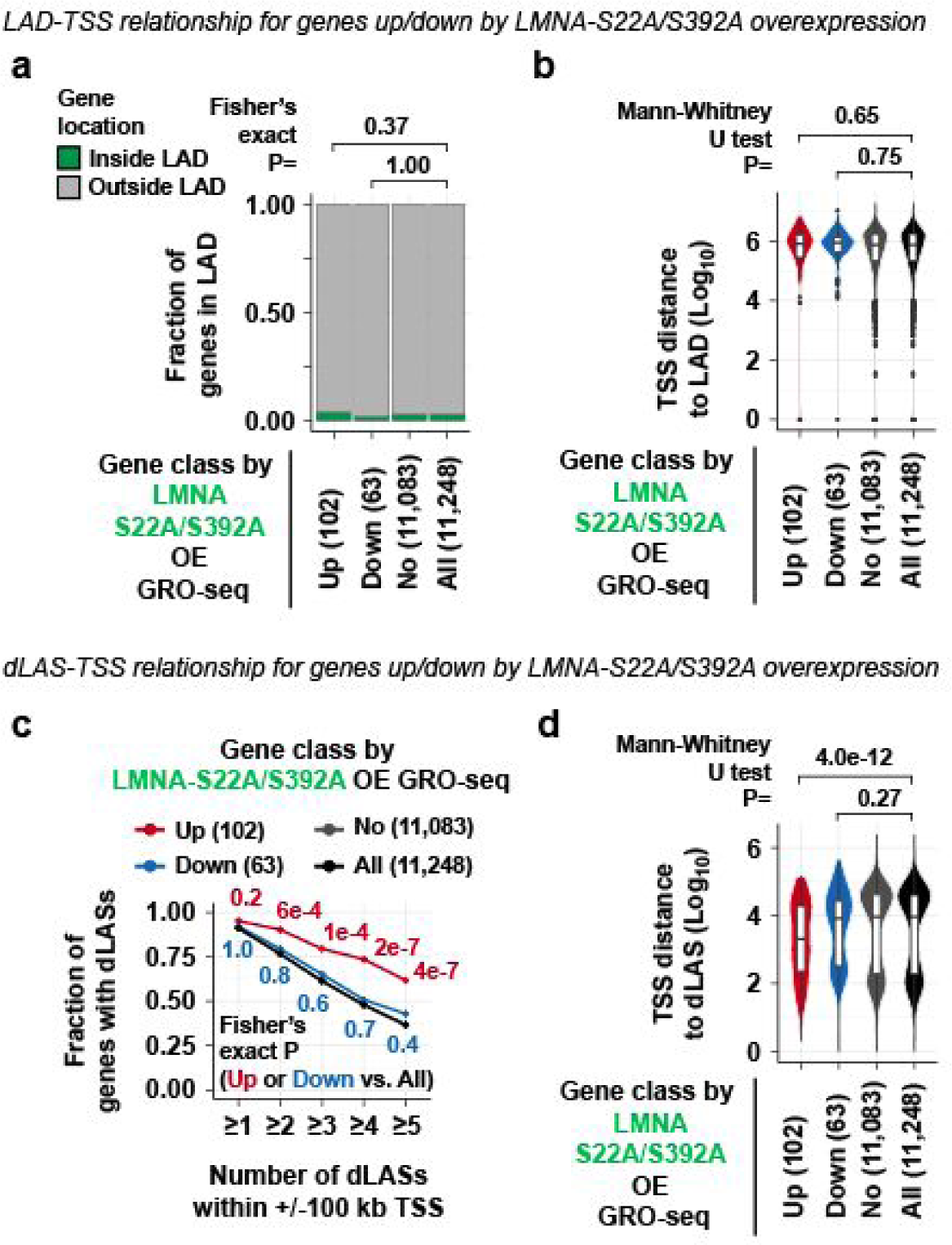
Overrepresentation of dLASs near genes de-repressed by LMNA-S22A/S392A overexpression. **a.** Fraction of genes whose TSSs are located inside or outside LADs. Genes are classified by the transcriptional changes in cells overexpressing (OE) the LMNA-S22A/S392A phospho-deficient mutant relative to wild-type BJ-5ta cells as determined by GRO-seq. **b.** TSS-to-LAD distance for genes differentially expressed in LMNA-S22A/S392A OE. **c.** Fraction of genes with dLASs within +/– 100 kb of TSS for genes differentially expressed in LMNA-S22A/S392A OE. Fraction is computed for each dLAS number cutoff indicated at the x-axis. **d.** TSS-to-dLASs distance for genes differentially expressed in LMNA-S22A/S392A OE.

**Extended Data Figure 4.**
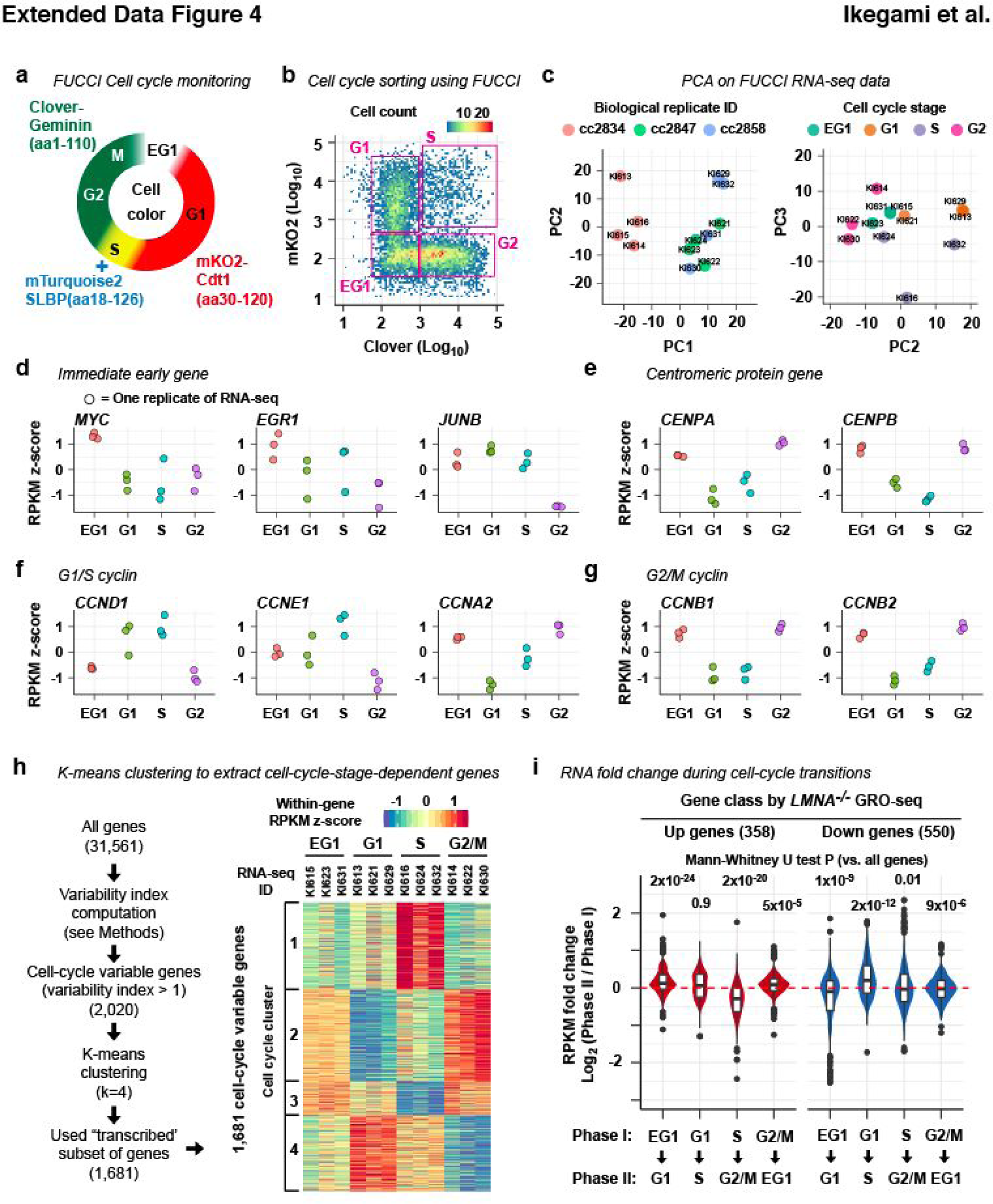
RNA-seq on FUCCI-sorted wild-type BJ-5ta cells. **a.** Schematic of the FUCCI experimental system. mKO2-Cdt1^30-120^ accumulates in the cell from G1 to S. Clover-Geminin^1-110^accumulates from S to M. mTurquoise2-SLBP^18-126^ accumulates in S. **b.** Sorting BJ-5ta cells expressing the FUCCI fluorescent proteins. **c.** PCA analysis on the FUCCI RNA-seq data set. Experiments were performed in triplicates. One biological replicate produces 4 samples, EG1, G1, S, and G2/M. PC1 distinguishes samples by biological replicates. The PC2 and 3 space distinguishes samples by cell-cycle stages. **d–g.** Known expression dynamics of representative cell-cycle-regulated genes demonstrated by the FUCCI RNA-seq dataset. For each gene, RPKM is normalized as z-scores within biological replicates. **h.** K-means clustering of genes by cell-cycle expression dynamics. (Left) Schematic of data analysis. Genes whose expression is variable between cell cycle stages are isolated and clustered by K-means clustering (k=4) (**Methods**). (Right) Each column represents each dataset of FUCCI-sorted RNA-seq. **i.** Log_2_ fold-change of expression between meighboring cell-cycle stages. Mann-Whitney test compares the distribution of log_2_ fold-change scores between *LMNA*^*–/–*^-up or *LMNA*^*–/–*^-down genes and all transcribed genes.

**Extended Data Figure 5.**
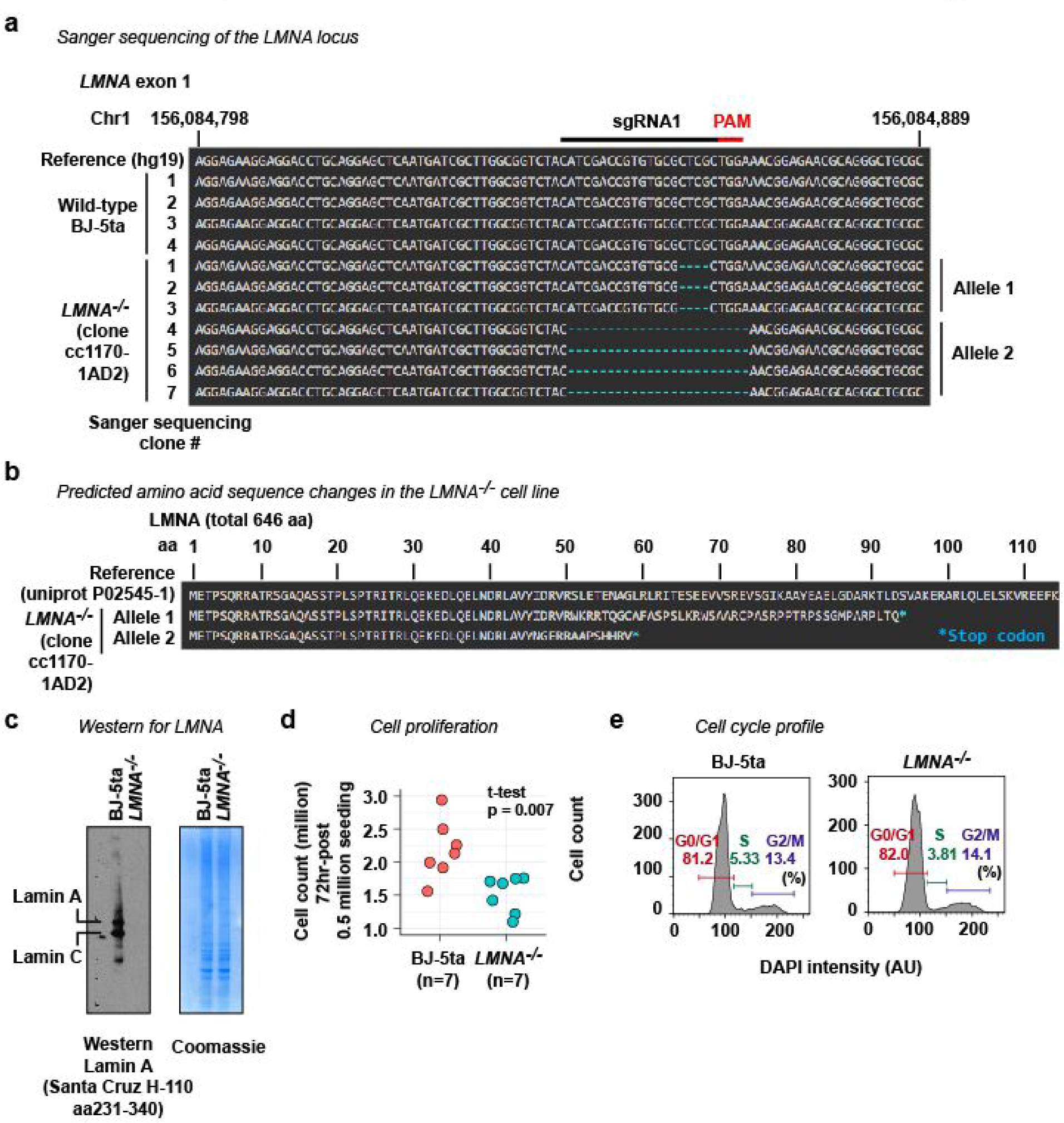
Characterization of *LMNA*^*–/–*^ cell line. **a.** The DNA sequence of the CRISPR-target site (sgRNA1) at the exon 1 of the *LMNA* gene. The *LMNA*^*–/–*^ cell line used in this study (clone cc1170-1AD2) is derived from BJ-5ta by CRISPR. Sanger sequencing revealed two modified alleles in the *LMNA*^*–/–*^ cell line. **b.** Both alleles in the *LMNA*^*–/–*^ cell line introduce premature stop codons. **c.** Western blotting for LMNA confirming loss of Lamin A and Lamin C, two products of the *LMNA* gene in the *LMNA*^*–/–*^ cell line. The lack of LMNA was also confirmed by immunofluorescence (**Fig. 1b**). **d.** The *LMNA*^*–/–*^ cell line proliferates slowly than the wild-type BJ-5ta cell line. **e.** Cell-cycle profiling by flow cytometry analysis of cellular DNA content. The *LMNA*^*–/–*^ cell line showed a cell-cycle profile comparable to that of wild-type BJ-5ta.ss

